# Stimulus-driven brain rhythms within the alpha band: The attentional-modulation conundrum

**DOI:** 10.1101/336941

**Authors:** Christian Keitel, Anne Keitel, Christopher SY Benwell, Christoph Daube, Gregor Thut, Joachim Gross

**Affiliations:** Institute of Neuroscience and Psychology, University of Glasgow, 62 Hillhead Street, Glasgow G12 8QB, UK; Psychology, School of Social Sciences, University of Dundee, Scrymgeour Building, Dundee DD1 4HN, UK; Institut für Biomagnetismus und Biosignalanalyse, Westfälische Wilhelms-Universität, Malmedyweg 15, 48149 Münster, Germany

**Keywords:** alpha rhythm, entrainment, phase synchronisation, spatial attention, steady-state response (SSR), frequency tagging

## Abstract

Two largely independent research lines use rhythmic sensory stimulation to study visual processing. Despite the use of strikingly similar experimental paradigms, they differ crucially in their notion of the stimulus-driven periodic brain responses: One regards them mostly as synchronised (entrained) intrinsic brain rhythms; the other assumes they are predominantly evoked responses (classically termed steady-state responses, or SSRs) that add to the ongoing brain activity. This conceptual difference can produce contradictory predictions about, and interpretations of, experimental outcomes. The effect of spatial attention on brain rhythms in the alpha-band (8 – 13 Hz) is one such instance: alpha-range SSRs have typically been found to *increase* in power when participants focus their spatial attention on laterally presented stimuli, in line with a gain control of the visual evoked response. In nearly identical experiments, retinotopic *decreases* in entrained alpha-band power have been reported, in line with the inhibitory function of intrinsic alpha. Here we reconcile these contradictory findings by showing that they result from a small but far-reaching difference between two common approaches to EEG spectral decomposition. In a new analysis of previously published human EEG data, recorded during bilateral rhythmic visual stimulation, we find the typical SSR gain effect when emphasising stimulus-locked neural activity and the typical retinotopic alpha suppression when focusing on ongoing rhythms. These opposite but parallel effects suggest that spatial attention may bias the neural processing of dynamic visual stimulation via two complementary neural mechanisms.

**SIGNIFICANCE STATEMENT:** Attending to a visual stimulus strengthens its representation in visual cortex and leads to a retinotopic suppression of spontaneous alpha rhythms. To further investigate this process, researchers often attempt to phase-lock, or entrain, alpha through rhythmic visual stimulation under the assumption that this entrained alpha retains the characteristics of spontaneous alpha. Instead, we show that the part of the brain response that is phase-locked to the visual stimulation *increased* with attention (in line with steady-state evoked potentials), while the typical suppression was only present in non-stimulus-locked alpha activity. The opposite signs of these effects suggest that attentional modulation of dynamic visual stimulation relies on two parallel cortical mechanisms – retinotopic alpha suppression and increased temporal tracking.

## INTRODUCTION

Cortical visual processing has long been studied using rhythmic sensory stimulation (Adrian and Matthews, 1934; Walter et al., 1946; Regan, 1966). This type of stimulation drives continuous brain responses termed steady-state responses (SSRs) that reflect the temporal periodicities in the stimulation precisely. SSRs allow tracking of individual stimuli in multi-element displays (Vialatte et al., 2010; Norcia et al., 2015). Further, they readily indicate cognitive biases of cortical visual processing, such as the selective allocation of attention (Morgan et al., 1996; Keitel et al., 2013; Stormer et al., 2014).

Although SSRs can be driven using a wide range of frequencies (Herrmann, 2001), stimulation at alpha band frequencies (8 – 13 Hz) has stirred particular interest. Alpha rhythms dominate brain activity in occipital visual cortices (Groppe et al., 2013; Keitel and Gross, 2016) and influence perception (Benwell et al., 2017; Iemi et al., 2017; Samaha et al., 2017; Benwell et al., 2018). Researchers have therefore used alpha-rhythmic visual stimulation in attempts to align the phase of – or *entrain –* intrinsic alpha rhythms and consequently provided evidence for visual alpha entrainment (Mathewson et al., 2012; Zauner et al., 2012; Spaak et al., 2014; Gulbinaite et al., 2017). These findings suggest that at least part of the SSR driven by alpha-band stimulation should be attributed to entrained alpha generators (Notbohm et al., 2016).

Some issues remain with such an account (Capilla et al., 2011; Keitel et al., 2014). For instance, experiments have consistently reported SSR power increases when probing effects of spatial selective attention on SSRs driven by lateralised hemifield stimuli (Müller et al., 1998a), also when using alpha-band frequencies (Kim et al., 2007; Kashiwase et al., 2012; Keitel et al., 2013). However, recent studies that used similar paradigms, but treated alpha-frequency SSRs as phase-entrained alpha rhythms in line with an earlier study using rhythmic transcranial magnetic stimulation (Herring et al., 2015), reported the opposite effect (Kizuk and Mathewson, 2017; Gulbinaite et al., 2019). Oscillatory brain activity showed attentional modulations characteristic of the intrinsic alpha rhythm during stimulation: Alpha power decreased over the hemisphere contralateral to the attended position, an effect known to be part of a retinotopic alpha power lateralisation during selective spatial attention (Worden et al., 2000; Kelly et al., 2006; Thut et al., 2006; Rihs et al., 2007; Capilla et al., 2012). Briefly put, studies analysing SSRs show a power *increase*, whereas studies analysing “entrained alpha” show a power *decrease* with attention.

Both neural responses originate from visual cortices contralateral to the hemifield position of the driving stimuli (Keitel et al., 2013; Spaak et al., 2014). Assuming a single underlying neural process, opposite attention effects therefore seemingly contradict each other. However, results in support of alpha entrainment differed in how exactly responses to the periodic stimulation were quantified. Effects consistent with SSR modulation resulted from spectral decompositions performed on trial-averaged EEG waveforms. This approach tunes the resulting power estimate to the part of the neural response that is sufficiently time-locked to the stimulation (Tallon-Baudry et al., 1996; Delorme and Makeig, 2004). Effects consistent with alpha entrainment instead typically result from averages of single-trial spectral transforms, thus emphasising intrinsic non-phase-locked activity (Tallon-Baudry et al., 1998; Herrmann et al., 2004). Both approaches have been applied before to compare stimulus-evoked and induced brain rhythms in alpha (Moratti et al., 2007) and gamma frequency ranges (~40 Hz; Tallon-Baudry et al., 1998; Picton et al., 2003). Here we focussed on contrasting the attentional modulation of alpha during- and SSRs driven by an alpha-rhythmic stimulation.

We therefore compared the outcome of both approaches in a new analysis of previously reported EEG data (Keitel et al., 2017b). Participants viewed two lateralised stimuli, both flickering at alpha band frequencies (10 and 12 Hz). They were cued to focus on one of the two and perform a target detection task at the attended position. We quantified spectral power estimates according to both approaches described above from the same EEG data. Should the outcome depend on the approach taken, we expected to find the typical alpha power lateralisation (contralateral < ipsilateral) when averaging single-trial power spectra. In power spectra of trial-averaged EEG instead we expected the typical SSR power gain modulation in the opposite direction (contralateral > ipsilateral). Crucially, such an outcome would warrant a re-evaluation of stimulus-driven brain rhythms in the alpha range and intrinsic alpha as a unitary phenomenon (alpha entrainment).

## METHODS

### Participants

For the present report, we re-analysed EEG data of 17 volunteers recorded in an earlier study (Keitel et al., 2017a). Participants (13 women; median age = 22 yrs, range = 19 – 32 yrs) declared normal or corrected-to-normal vision and no history of neurological diseases or injury. All procedures were approved by the ethics committee of the College of Science & Engineering at the University of Glasgow (application no. 300140020) and adhered to the guidelines for the treatment of human subjects in the Declaration of Helsinki. Volunteers received monetary compensation of £6/h. They gave informed written consent before participating in the experiment. Note that we excluded five additional datasets on grounds reported in the original study (four showed excessive eye movements, one underperformed in the task).

### Stimulation

Participants viewed experimental stimuli on a computer screen (refresh rate = 100 frames per sec) at a distance of 0.8 m that displayed a grey background (luminance = 6.5 cd/m^2^). Small concentric circles in the centre of the screen served as a fixation point (*Figure 1*). Two blurry checkerboard patches (horizontal/vertical diameter = 4° of visual angle) were positioned at an eccentricity of 4.4° from central fixation, one each in the lower left and lower right visual quadrants. Both patches changed contrast rhythmically during trials: Stimulus contrast against the background was modulated by varying patch peak luminance between 7.5 cd/m^2^ (minimum) and 29.1 cd/m^2^ (maximum).

**Figure 1.**
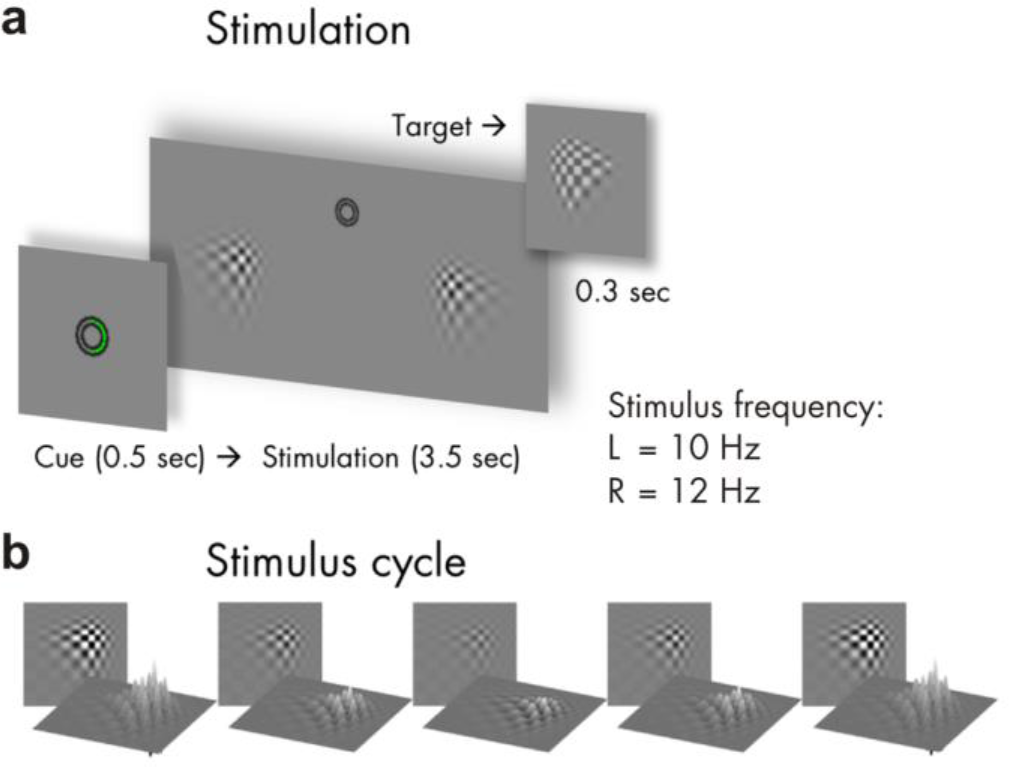
Stimulus schematics and trial time course. (**a**) shows the time course of one trial with a cue displayed for 0.5 sec (here: Attend Right), followed by the bilateral visual stimulation for 3.5 sec. Left (L) stimulus contrast fluctuated with a rate of 10 Hz and Right (R) stimulus contrast at 12 Hz. Targets that participants were instructed to respond to were slightly altered versions of the stimuli (see inset) that were displayed occasionally for 0.3 sec. (**b**) Rhythmic visual stimulation was achieved by a frame-by-frame adjustment of global stimulus contrast (through local luminance changes) as exemplified here in one representative cycle.

On each screen refresh, peak luminance changed incrementally to approach temporally smooth contrast modulations as opposed to a simple on-off flicker (Andersen and Muller, 2015). Further details of the stimulation can be found in Keitel et al. (2017a). The contrast modulation followed a 10-Hz periodicity for the left and a 12-Hz periodicity for the right stimulus. Note that the experiment featured further conditions displaying quasi-rhythmic contrast modulations in different frequency bands. Corresponding results can be found in the original report and will not be considered in the present analysis.

### Procedure and Task

Participants performed the experiment in an acoustically dampened and electromagnetically shielded chamber. In total, they were presented with 576 experimental trials, subdivided into 8 blocks with durations of ~5 min each. Between blocks, participants took self-paced breaks. Prior to the experiment, participants practiced the behavioural task (see below) for at least one block. After each block they received feedback regarding their accuracy and response speed. The experiment was comprised of 8 conditions (= 72 trials each) resulting from a manipulation of the two factors attended position (left vs. right patch) and stimulation frequency (one rhythmic and three quasi-rhythmic conditions) in a fully balanced design. Trials of different conditions were presented in pseudo-random order. As stated above, the present study focussed on the two conditions featuring fully rhythmic stimuli. Corresponding trials (N = 144) were thus selected a posteriori from the full design.

Single trials began with cueing participants to attend to the left or right stimulus for 0.5 sec, followed by presentation of the dynamically contrast-modulating patches for 3.5 sec (*Figure 1*). After patch offset, an idle period of 0.7 sec allowed participants to blink before the next trial started.

To control whether participants maintained a focus of spatial attention, they were instructed to respond to occasional brief “flashes” (0.3 sec) of the cued stimulus (= targets) while ignoring similar events in the other stimulus (= distracters). Targets and distracters occurred in one third of all trials and up to 2 times in one trial with a minimum interval of 0.8 sec between subsequent onsets. Detection was reported as speeded responses to flashes (recorded as space bar presses on a standard keyboard).

### Behavioural data recording and analyses

Flash detections were considered a ‘hit’ when a response occurred from 0.2 to 1 sec after target onset. Delays between target onsets and responses were considered reaction times (RT). Statistical comparisons of mean accuracies (proportion of correct responses to the total number of targets and distracters) and median RTs between experimental conditions were conducted and reported in (2017a). In the present study, we did not consider the behavioural data further. Note that the original statistical analysis found that task performance in Attend-Left and Attend-Right conditions was comparable.

### Electrophysiological data recording

EEG was recorded from 128 scalp electrodes and digitally sampled at a rate of 512 Hz using a BioSemi ActiveTwo system (BioSemi, Amsterdam, Netherlands). Scalp electrodes were mounted in an elastic cap and positioned according to an extended 10-20-system (Oostenveld and Praamstra, 2001). Lateral eye movements were monitored with a bipolar outer canthus montage (horizontal electro-oculogram). Vertical eye movements and blinks were monitored with a bipolar montage of electrodes positioned below and above the right eye (vertical electro-oculogram).

### Electrophysiological data pre-processing

From continuous data, we extracted epochs of 5 s, starting 1 s before patch onset using the MATLAB toolbox EEGLAB (Delorme and Makeig, 2004). In further pre-processing, we excluded epochs that corresponded to trials containing transient targets and distracters (24 per condition) as well as epochs with horizontal and vertical eye movements exceeding 20 μV (~ 2° of visual angle) or containing blinks. For treating additional artefacts, such as single noisy electrodes, we applied the ‘fully automated statistical thresholding for EEG artefact rejection’ (FASTER; Nolan et al., 2010). This procedure corrected or discarded epochs with residual artefacts based on statistical parameters of the data. Artefact correction employed a spherical-spline-based channel interpolation. Epochs with more than 12 artefact-contaminated electrodes were excluded from analysis.

From 48 available epochs per condition, we discarded a median of 14 epochs for the Attend-Left conditions and 15 epochs for the Attend-Right conditions per participant with a between-subject range of 6 to 28 (Attend-Left) and 8 to 31 epochs (Attend-Right). Within-subject variation of number of epochs per condition remained small with a median difference of 3 trials (maximum difference = 9 for one participant).

Subsequent analyses were carried out in Fieldtrip (Oostenveld et al., 2011) in combination with custom-written routines. We extracted segments of 3 s starting 0.5 s after patch onset from pre-processed artefact-free epochs (5 s). Data prior to stimulation onset (1 s), only serving to identify eye movements shortly before and during cue presentation, were omitted. To attenuate the influence of stimulus-onset evoked activity on EEG spectral decomposition, the initial 0.5 s of stimulation were excluded. Lastly, because stimulation ceased after 3.5 s, we also discarded the final 0.5 s of original epochs.

### Electrophysiological data analyses – spectral decomposition

Artefact-free 3-sec epochs were converted to scalp current densities (SCDs), a reference-free measure of brain electrical activity (Ferree, 2006; Kayser and Tenke, 2015), by means of the spherical spline method (Perrin et al., 1987) as implemented in Fieldtrip (function *ft_scalpcurrentdensity*, method ‘spline’, lambda = 10^−4^). Detrended (i.e. mean and linear trend removed) SCD time series were then Tukey-tapered and subjected to Fourier transforms while employing zero-padding in order to achieve a frequency-resolution of 0.25 Hz. Crucially, from resulting complex Fourier spectra we calculated two sets of aggregate power spectra with slightly different approaches. First, we calculated power spectra as the average of squared absolute values of complex Fourier spectra (Z) as follows:

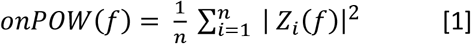

where *onPOW* is the classical power estimate for ongoing (intrinsic) oscillatory activity for frequency *f* and *n* is the number of trials. Secondly, we additionally calculated the squared absolute value of the averaged complex Fourier spectra according to:

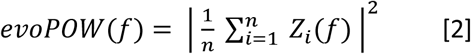

The formula yields *evoPOW*, or evoked power, an estimate that is identical with the frequency-tagging standard approach of averaging per-trial EEG time series before spectral decomposition. This step is usually performed to retain only the truly phase-locked response to the stimulus (Tallon-Baudry et al., 1996). Note that both formulas only differ in the order in which weighted sums and absolute values are computed. Also note that formula [2] is highly similar to the calculation of intertrial phase coherence (ITC), a popular measure of phase locking (Cohen, 2014; Gross, 2014; van Diepen and Mazaheri, 2018). ITC calculation additionally includes a trial-by-trial amplitude normalisation. To complement our analysis we thus quantified ITC according to:

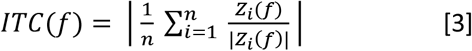

For further analyses, power spectra were normalised by converting them to decibel scale, i.e. taking the decadic logarithm, then multiplying by 10 (hereafter termed log power spectra). ITC was converted to ITCz to reduce the bias introduced by differences in trial numbers between conditions (Bonnefond and Jensen, 2012; Samaha et al., 2015).

### Alpha power – attentional modulation and lateralisation

Spectra of ongoing power (*onPOW*), pooled over both experimental conditions and all electrodes, showed a prominent peak in the alpha frequency range (*Figure 2*). We used mean log ongoing power across the range of 8 – 13 Hz to assess intrinsic alpha power modulations by attention. Analysing Attend-Right and Attend-Left conditions separately, yielded two alpha power topographies for each participant. These were compared by means of cluster-based permutation statistics (Maris and Oostenveld, 2007) using *N* = 5000 random permutations. We clustered data across channel neighbourhoods with an average size of 7.9 channels that were determined by triangulated sensor proximity (function *ft_prepare_neighbours*, method ‘triangulation’). The resulting probabilities (*P*-values) were corrected for two-sided testing. Subtracting left-lateralised (Attend-Left conditions) from right-lateralised (Attend-Right) alpha power topographies, we found a right-hemispheric positive and a left-hemispheric negative cluster of electrodes that was due to the retinotopic effects of spatial attention on alpha power lateralisation (*Figure 3*), similar to an earlier re-analysis of the other conditions of this experiment (Keitel et al., 2018).

**Figure 2.**
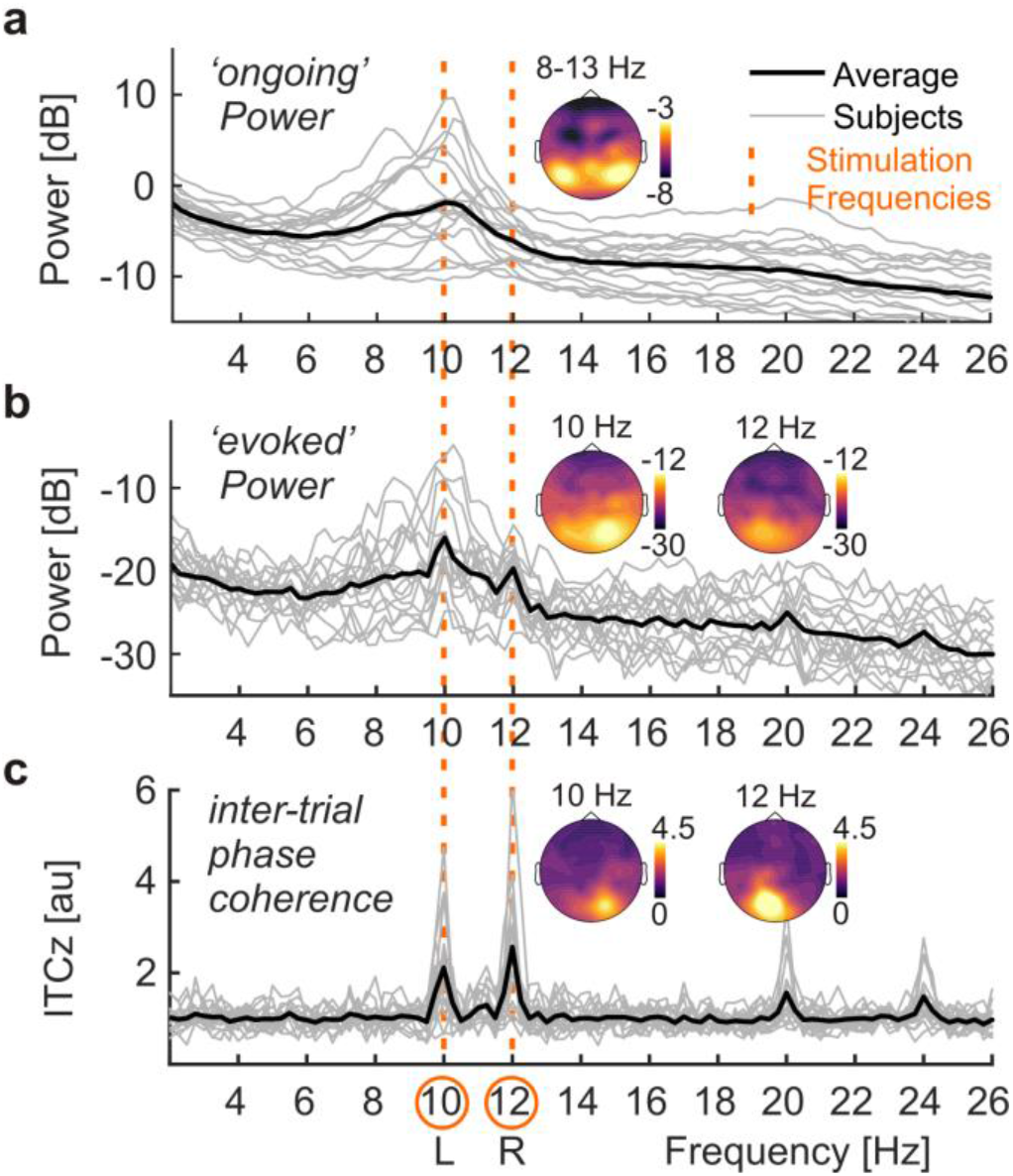
EEG spectral decomposition. (**a**) Power spectra collapsed across conditions and all electrode positions below the sagittal midline for single subjects (light grey lines) and group averages (strong black line). Note the characteristic alpha peaks in the frequency range of 8 – 13 Hz. Inset scalp map shows topographical distribution of alpha power on a dB scale based on scalp current densities. (**b**) Same as in (a) but for ‘evoked’ power. Distinct peaks are visible at stimulation frequencies 10 & 12 Hz (dashed vertical orange lines across plots). Inset scalp maps show topographical distributions of SSR power at 10 & 12 on a dB scale. Note the difference in scale between ongoing power in (a) and evoked power (b). (**c**) Same as in (a) but for inter-trial phase-locking (ITCz). Inset scalp maps show topographical distributions of SSR ITCz at 10 & 12.

**Figure 3.**
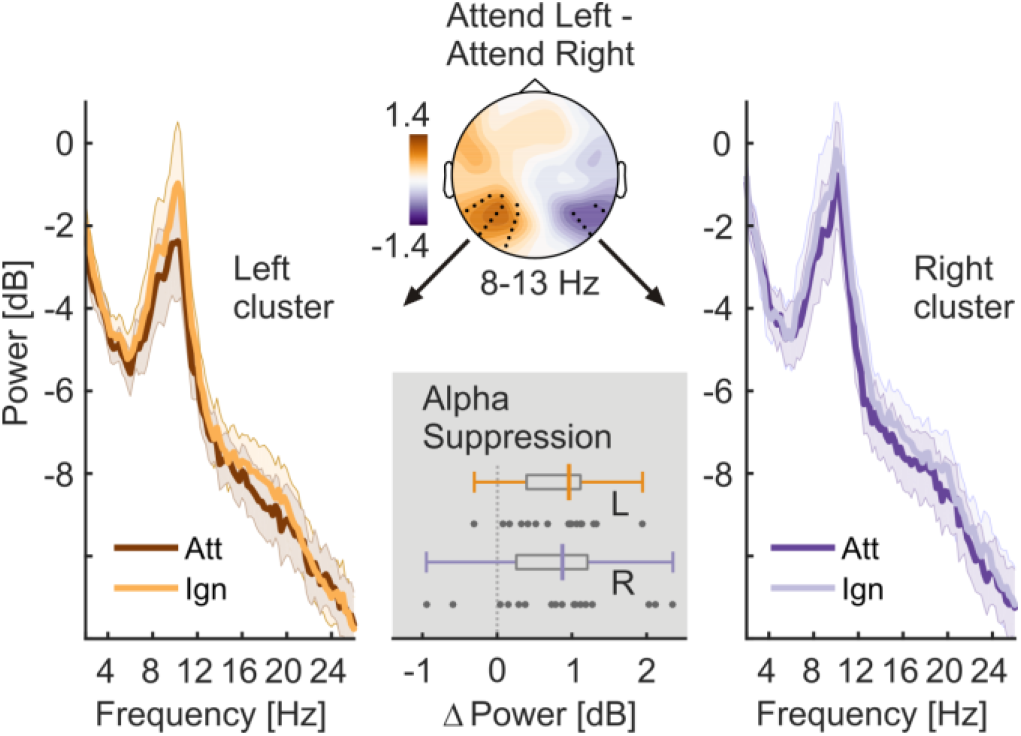
Allocation of spatial attention produces retinotopic alpha power modulation. The scalp map (top, center) depicts alpha power lateralisation (Attend Left – Attend right conditions) on a dB scale. Black dots indicate left- and right-hemispheric electrode-clusters that showed a consistent difference in group statistics (two-tailed cluster-based permutation tests). Left and right spectra illustrate alpha power differences in respective clusters when the contralateral hemifield was attended (Att) versus ignored (Ign). The bottom grey inset depicts the distribution of individual alpha power suppression effects (Ignored minus Attended) within left (L) and right (R) hemisphere clusters in the 8 – 13 Hz band. Boxplots indicate interquartile ranges (boxes) and medians (coloured vertical intersectors). Dots below show individual effects (1 dot = 1 participant).

Finally, we tested the difference between Attend-Left and Attend-Right conditions, i.e. attention effects for left- and right-hemispheric clusters separately. To this end, we submitted alpha power differences (contralateral hemifield attended minus ignored) to Bayesian one-sample t-tests against zero (Rouder et al., 2009). Attention effects were further compared against each other by means of a Bayesian paired-samples t-test as implemented in JASP (JASP-Team, 2018) with a Cauchy prior scaled to r = 0.5, putting more emphasis on smaller effects (Rouder et al., 2012; Schonbrodt and Wagenmakers, 2017).

This procedure allowed us to quantify the evidence in favour of the null vs the alternative hypothesis (H_0_ vs H_1_). For each test, the corresponding Bayes factor (called BF_10_) showed evidence for H_1_ (compared to H_0_) if it exceeded a value of 3, and no evidence for H_1_ if BF_10_ < 1, with the intervening range 1 – 3 termed ‘anecdotal evidence’ by convention (Wagenmakers et al., 2011). Inversing BF_10_, to yield a quantity termed BF_01_, served to quantify evidence in favour of H_0_ on the same scale. For BF_10_ and BF_01_, values < 1 were taken as inconclusive evidence for either hypothesis. Note that for the sake of brevity we report errors in BF estimates only when exceeding 0.001%.

### SSR power – attentional modulation

Spectra of evoked power, pooled over both experimental conditions and all electrodes, revealed periodic responses to the two stimuli at the respective stimulation frequencies, 10 and 12 Hz (*Figure 2*). Therefore, we assessed attention effects for these two spectral SSR representations. Two separate cluster-based permutation tests, one for each stimulation frequency, contrasted evoked power topographies between attended and ignored (= other stimulus attended) conditions. Two-sided tests were performed with the same parameters as for alpha power (see above).

Again, we found one electrode cluster carrying systematic attention effects per frequency. As for alpha, SSR power from these two clusters were subjected to separate Bayesian one-sample t-tests against zero (one-sided, attended > ignored) and compared against each other by means of a Bayesian paired-sample t-test (two-sided).

### SSR inter-trial phase coherence – attentional modulation

We also evaluated a pure measure of neural phase-locking to the stimulation, SSR inter-trial phase coherence (ITC), because evoked power can be regarded as a hybrid measure depending on both the amplitude of the underlying rhythmic response and the consistency of its phase across trials. ITC indicates only the latter as SSRs are set to unit amplitude prior to summing across trials (see formula 3). ITC spectra, pooled over both experimental conditions and all electrodes, showed distinct neural phase-locking at the respective driving frequencies, 10 and 12 Hz (*Figure 2*). Cluster-based permutation testing confirmed topographic regions that showed systemic gain effects in ITC. Subsequently, the same Bayesian inference was applied to data from these clusters as for SSR power.

### Correlation of alpha and SSR attention effects – group level

As a consequence of our counter-intuitive finding that SSR attention effects appeared strongest over occipital regions ipsi-lateral to the driving stimulus (see Results section *SSR power & inter-trial phase locking – attentional modulation* below), we explored a posteriori whether these effects could be explained by ipsilateral increases in alpha power during focussed attention. We correlated attention effects on alpha and SSR power using Bayesian inference (rank correlation coefficient Kendall’s tau-b or *τ_b_*, beta-prior = 0.75) to test for a positive linear relationship. More specifically, we correlated the left-hemispheric alpha power suppression (Ignored minus Attended) with the 10-Hz SSR (evoked) power attention effect (Attended minus Ignored) and the right-hemispheric alpha power suppression with the 12-Hz SSR power attention effect. We opted for these combinations because the corresponding effects overlapped topographically (see Results). Along with the correlation coefficient ρ, we report its 95%-Credible Interval (95%-CrI).

We also probed the linear relationship between alpha power and SSR ITC attention effects. Because ITC gains were not clearly lateralised, we collapsed gain effects (Attended minus Ignored) across both stimulation frequencies and correlated these with a hemisphere-collapsed alpha suppression index. This index was quantified as the halved sum of left and right-hemispheric suppression effects as retrieved from significant clusters in the topographical analysis of alpha power differences (Attend Left minus Attend Right), shown in *Figure 3*. Again, we expected a positive correlation here if alpha power suppression influenced phase-locking to visual stimulation. For means of comparison, we repeated this analysis with attention effects on SSR power collapsed across frequencies.

### Alpha and SSR attention effects – subject level regression

The relationship between alpha power (lateralisation) and SSR attentional modulation was further subjected to a more fine-grained analysis considering within-subject variability across single trials and allowing for a better control of between-subject differences in alpha and SSR power. We assumed that if the SSR attention effect (i.e. the ipsilateral SSR power gain) was a mere consequence of the co-localised alpha power increase then these two effects should co-vary across trials. For this analysis we recalculated single-trial alpha power and SSR evoked power / ITC estimates at each EEG sensor and for both conditions in each subject based on the same artefact-removed EEG epochs and using the same spectral decomposition as described above. Because ITC is not defined for single trials, we used a Jackknife approach that computed single trial estimates in a leave-one-out procedure and allowed for subsequent evaluation of inter-trial variability (Richter et al., 2015). For consistency, we computed similar alpha-power Jackknife estimates. From these estimates, we calculated attention effects as all possible pairwise differences between trials of different conditions (Attend Left vs Attend Right), yielding distributions of alpha power hemispheric lateralisation and SSR evoked power / ITC attentional modulation (for 10 & 12 Hz SSRs separately). To validate this approach, we used it to reproduce alpha power and SSR attention effects described below (data not shown, reproducible via code in online repository (Keitel et al., 2017b)).

We then tested for a linear relationship between both z-scored measures by subjecting them to a robust linear regression (MATLAB function ‘robustfit’, default options), carried out for each EEG sensor separately. The obtained subject-specific regression coefficients *β* (slopes) were entered into a group statistical test. We tested slopes against zero (i.e. no linear relationship) by means of cluster-based permutation tests (two-tailed), clustering across EEG sensors. Four tests were carried out in total; one for each regression of alpha power lateralisation with SSR evoked power or SSR ITC attentional modulation, and separately for 10 & 12 Hz SSR, respectively. This procedure was supplemented by sensor-by-sensor Bayesian t-tests (Rouder et al., 2009) to quantify the evidence in favour of a linear vs no relationship (see Methods section *Alpha power – attentional modulation and lateralisation* regarding Bayesian inference).

## RESULTS

### Ongoing alpha power – attentional modulation and lateralisation

The power of the ongoing alpha rhythm lateralised with the allocation of spatial attention to left and right stimuli. A topographic map of the differences in alpha power between Attend-Left and Attend-Right conditions shows significant left- and right-hemispheric electrode clusters (*Figure 3*). These clusters signify retinotopic alpha power modulation when participants attended to left vs right stimulus positions (right cluster: t_sum_ = −21.454, *P* = 0.026; left cluster: t_sum_ = 81.264, *P* = 0.002). The differences are further illustrated in power spectra pooled over electrodes of each cluster (*Figure 3*). As predicted, alpha power at each cluster was lower when participants attended to the contralateral stimulus. Bayesian inference confirmed the alpha power attention effect for the right (*M* = 0.806 dB, *SEM* = 0.216; *BF*_10_ = 21.17) and left cluster (*M* = 0.790 dB, *SEM* = 0.133; *BF*_10_ = 906.36). Both effects were of comparable magnitude (*BF*_01_ = 4.009 ± 0.007).

### SSR power & inter-trial phase locking – attentional modulation

Crucially, we found the opposite pattern when looking at SSRs, i.e. the exact same data but with a slightly different focus on oscillatory brain activity that was time-locked to the stimulation (compare formulas 1 and 2): SSRs showed increased power when the respective driving stimulus was attended versus ignored (*Figure 4*). The power of neural responses evoked by our stimuli (SSRs) was at least one order of magnitude smaller than ongoing alpha power on average (difference > 10dB, i.e. between 10 – 100 times). Nevertheless, SSRs could be clearly identified as distinct peaks in (evoked) power and ITC spectra. Consistent with the retinotopic projection to early visual cortices, topographical distributions of both measures showed a focal maxima contra-lateral to the respective stimulus positions that were attended (*Figure 2*). Counter-intuitively though, maximum attention effects on SSR power did not coincide topographically with sites that showed maximum SSR power overall (compare scalp maps in *Figure 2 & 4*). Also, due to their rather ipsilateral scalp distributions (with respect to the attended location), SSR attention effects did not match topographies of attention-related decreases in ongoing alpha power (compare scalp maps in *Figures 3 & 4*). The 10-Hz SSR driven by the left-hemifield stimulus showed a left-hemispheric power increase when attended (t_sum_ = 15.837, *P* = 0.059). Similarly, attention increased the power of the 12-Hz SSR driven by the right-hemifield stimulus in a right-hemispheric cluster (t_sum_ = 53.282, *P* < 0.001). Bayesian inference confirmed the attention effect on 10-Hz (*M* = 3.727 dB, *SEM* = 0.919; *BF*_10_ = 37.05) and 12-Hz SSR power (*M* = 4.473 dB, *SEM* = 0.841; *BF*_10_ = 329.75) averaged within clusters. Both effects were of comparable magnitude (*BF*_01_ = 3.443 ± 0.005).

**Figure 4.**
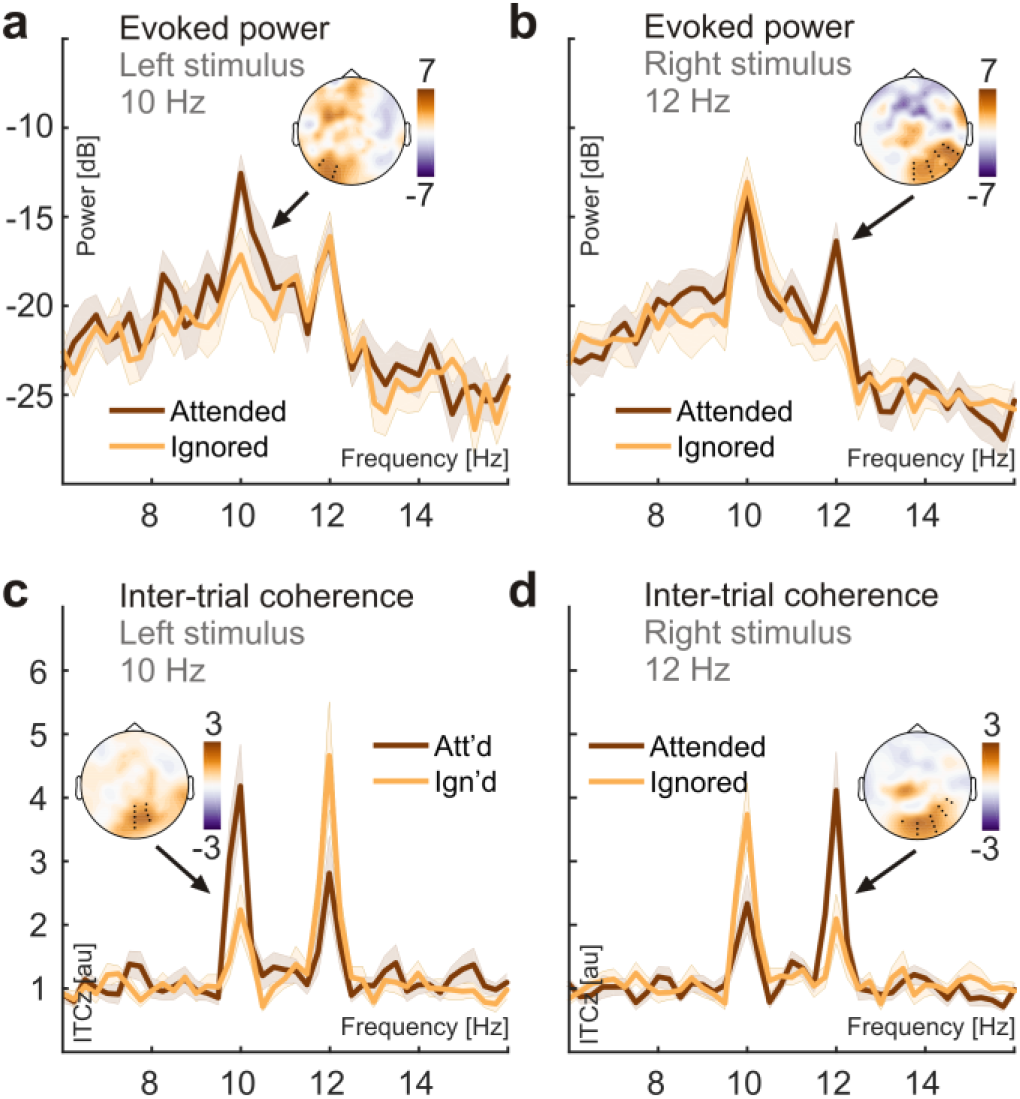
Attention effects on SSR evoked power (evoPow) and SSR inter-trial phase coherence. (**a**) SSR evoked power spectra show systematic power differences at the presentation frequency (10 Hz) of the left stimulus when it was attended (dark red) versus ignored (orange). The inset scalp map illustrates the topographical distribution of the attention effects. Power spectra were averaged across electrodes (black dots in scalp maps) that showed consistent attention effects in group statistics (two-tailed cluster-based permutation tests) for Attended and Ignored conditions separately. (**b**) Same as in (a) but for the 12-Hz stimulus presented in the right visual hemifield. (**c,d**) Same as in (a,b) but using ITCz as a measure of SSR inter-trial phase coherence.

SSR phase-locking (quantified as ITCz) also increased with attention to the respective stimulus. In contrast to evoked power, topographical representations of these effects showed greater overlap with the sites that showed maximum phase-locking in general (*Figure 4*). For both frequencies, ITCz increased in central occipital clusters (10 Hz: t_sum_ = 41.351, *P* = 0.004; 12 Hz: t_sum_ = 31.116, *P* = 0.012). Again, Bayesian inference confirmed the attention effect on 10-Hz (M = 1.386 au, *SEM* = 0.297; *BF*_10_ = 105.71, one-sided) and 12-Hz ITCz (*M* = 1.824 au, *SEM* = 0.451; *BF*_10_ = 36.11, one-sided). Evidence for a greater attention effect on 12-Hz than on 10-Hz ITC remained inconclusive (*B*_F10_ = 0.473).

### Correlation of alpha and SSR attention effects – group level

Lastly, we tested whether the SSR attention gain effects were mere reflections of the topographically coinciding ipsilateral ongoing alpha power increase during focussed attention that co-occurred with the contralateral ongoing alpha-power decrease (*Figure 3*). Speaking against this account, Bayesian inference provided moderate evidence against the expected positive correlations between the left-hemispheric alpha attention effect and the 10-Hz SSR attention effect (*τ_b_* = −0.221, 95%-CrI = [0.002 0.269]; BF_01_ = 5.811) and between the right-hemispheric alpha attention effect and the 12-Hz SSR attention effect (*τ_b_* = −0.088, 95%-CrI = [0.004 0.315]; BF_01_ = 3.904). These relationships are further illustrated by corresponding linear fits in *Figure 5*.

**Figure 5.**
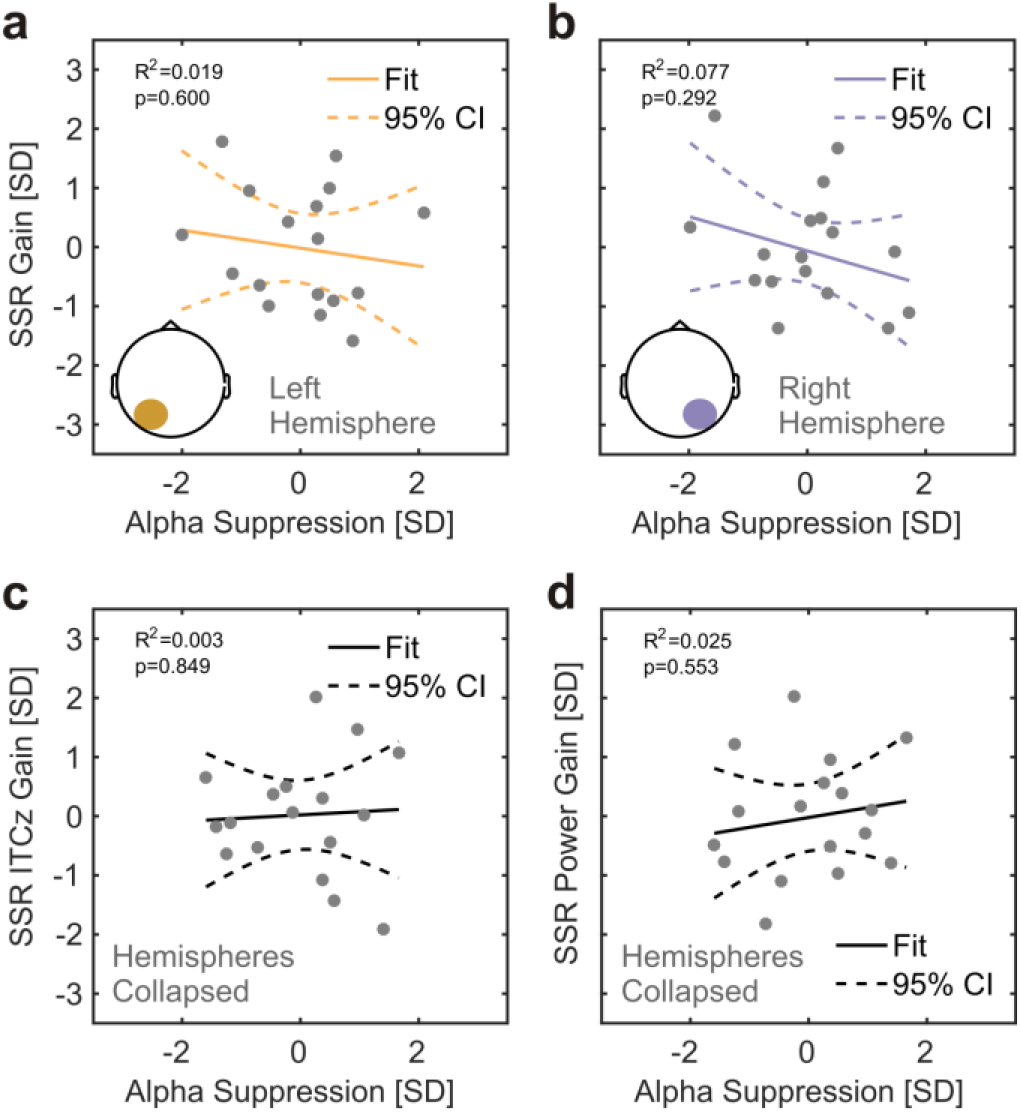
Relationships between attention effects on alpha power and SSRs. (**a**) Individual 10-Hz (left stimulus) SSR evoked power gain (Attended minus Ignored; z-scored, y-axis) as a function of alpha suppression (Ignored minus Attended; z-scored, x axis) in overlapping left-hemispheric parieto-occipital electrode clusters. Grey dots represent participants. Coloured lines depict a straight line fit and its confidence interval (dashed lines). Goodness of fit of the linear model provided as R^2^ along with corresponding P-Value. As confirmed by additional tests, both attention effects do not show a positive linear relationship that would be expected if the ipsilateral SSR power gain effect was a consequence of the ipsilateral alpha suppression. (**b**) Same as in (a), but for the 12 Hz SSR driven by the right stimulus in overlapping right-hemispheric parieto-occipital electrode clusters. (**c,d**) Similar to (a) but for attention-related gain effects on SSR ITCz (z-scored, y-axis) in (c) and gain effects on SSR evoked power in (d), both collapsed across electrode clusters showing 10- and 12-Hz SSR attention effects. Alpha suppression was collapsed across left- and right-hemispheric electrode clusters (see Figure 3).

Following this analysis, we further explored the relationship between spatially non-overlapping decreases in alpha-power contralateral to the attended position and the ipsilateral SSR power gain effects. For the lack of a specific hypothesis about the sign of the correlation in this case, we quantified the evidence for any relationship (two-sided test). The results remained inconclusive for a correlation between the left-hemispheric alpha attention effect and the right-hemispheric 12-Hz SSR attention effect (*τ_b_* = 0.235, 95%-CrI = [−0.110 0.487]; BF_01_ = 1.280) and between the right-hemispheric alpha attention effect and the left-hemispheric 10-Hz SSR attention effect (*τ_b_* = 0.103, 95%-CrI = [−0.218 0.383]; BF_01_ = 2.400).

Finally, we repeated this analysis for attention effects on inter-trial phase coherence (ITC). Because SSR ITC attention effects did not show a clear topographical lateralisation (*Figure 4*), they were collapsed across driving frequencies (10 & 12 Hz). Again, findings were inconclusive when looking into the correlation between these aggregate SSR ITC gain effects and a hemisphere-collapsed alpha suppression index (*τ_b_* = −0.059, 95%-CrI = [−0.349 0.251]; BF_01_ = 2.653). Correlating collapsed attention effects of SSR evoked power with the same pooled alpha suppression index yielded identical results regarding the rank correlation (also see linear fits in *Figure 5*).

### Alpha and SSR attention effects – subject level regression

A more fine-grained analysis of single-trial co-variation of alpha power lateralisation and SSR gain effects during focussed spatial attention largely corroborated the group level results. Clustering across EEG sensors, we found that only the 12-Hz SSR evoked power attention effect and alpha lateralisation co-varied systematically across participants at occipital sites (permutation test, *T_sum_* = −17.517, *p* = 0.023). The negative sign of the slope however contradicted the expected positive relationship (*Figure 6a*). Neither 10-Hz SSR evoked power nor SSR ITC (both frequencies) revealed similar systematic relationships with alpha power.

**Figure 6.**
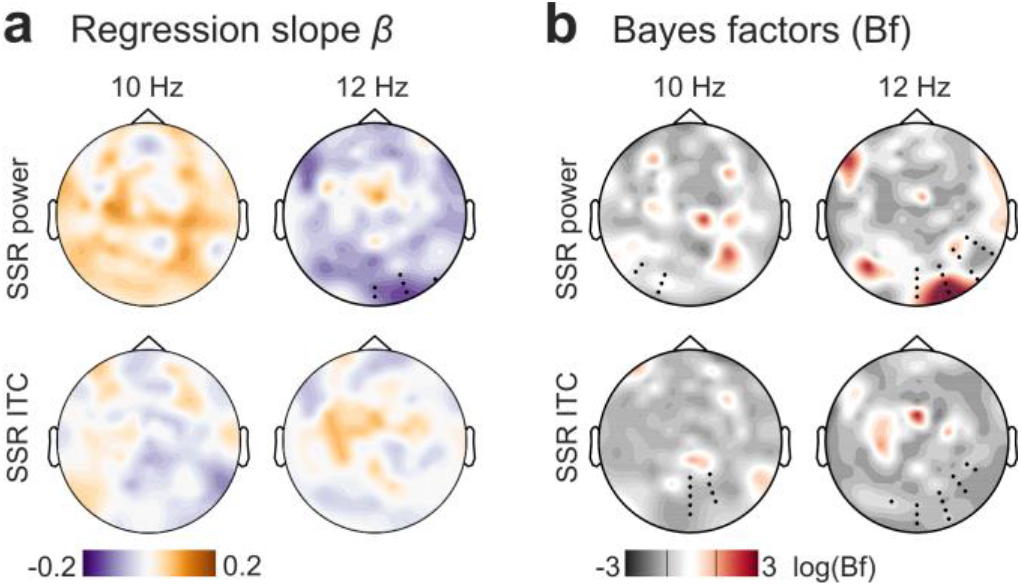
Summary of subject-level analysis of the linear relationship between alpha power and SSR attentional modulation. (**a**) depicts the topographical distribution of group-averaged (N=17) regression coefficients β (slopes) for SSR evoked power (top row) and SSR inter-trial coherence (ITC, bottom row), separated by SSR frequencies 10 Hz (left column) and 12 Hz (right column). Hot colours indicate a positive linear relationship and cool colours a negative relationship. Black dots in the upper right panel indicate a cluster of electrodes showing a systematic effect (p < 0.05, cluster-based permutation test) absent in tests illustrated in the other 3 panels. (**b**) Results of sensor-by-sensor group-level Bayesian inference (Bayesian t-tests) of regression slopes against zero, plotted as topographies on a log(BF_10_) scale. Plots arranged as in (a). Red colours indicate stronger evidence for H_1_, grey colours indicate stronger evidence for H_0_. Black lines in the colour scale below scalp maps denote thresholds that signal moderate evidence for H_0_ (log(1/3) = −1.099) or H_1_ (log(3) = −1.099) by convention. Superimposed black dots indicate clusters showing systematic attention effects on SSR evoked power / ITC as depicted in *Figure 4* for comparison.

Additionally, we used Bayesian inference on the distributions of individual regression slopes (indicating the linear relationship between alpha and SSR attention effects) by sensor to quantify the plausibility of either H_1_ or H_0_ in scalp maps (*Figure 6b*). We further overlaid these scalp maps with electrode clusters showing SSR attention effects (compare with *Figure 4*). Average Bayes factors (Bfs) within clusters indicated that evidence for or against any type of linear relationship remained inconclusive for 10-Hz (mean Bf_01_ = 1.422 range = 0.639 – 2.343) and 12-Hz SSR evoked power (mean Bf_01_ = 1.245 range = 0.153 – 3.673), although it should be mentioned that the 12-Hz cluster contained a local maximum (Bf_10_ = 1/Bf_01_ = 6.534) that coincided topographically with the effect identified by the cluster-based permutation test. For ITC evidence favoured H_0_, i.e. the absence of any relationship with 10-Hz (mean Bf_01_ = 3.040, range = 1.861 – 4.014) and 12-Hz SSR (Bf_01_ = 3.030, range = 1.391 – 4.016) was 3 times more likely given our data.

Our findings show a fine distinction between SSR evoked power and ITC gain effects with respect to a possible connection to alpha lateralisation in that only the latter provided conclusive evidence against such a relationship. As a likely explanation, SSR evoked power still contains residual alpha activity that confounds tests for covariation. Conversely, the single-trial power normalisation step undertaken during the calculation of SSR ITC makes it less susceptible to this confound. Taken together, the findings of this analysis do not support a positive linear relationship of alpha lateralisation and SSR gain effects (especially on ITC). Therefore, it is unlikely that the counter-intuitive topography of SSR attentional modulation is a reflection of alpha power lateralisation during focused spatial attention.

## DISCUSSION

We found that two common spectral measures of alpha-band EEG during alpha-rhythmic visual stimulation reflect effects of spatial attention with opposite signs. In the following we discuss how this finding supports the notion of two complementary neural mechanisms governing the cortical processing of dynamic visual input.

### Analysis approach determines sign of attentional modulation

When focussing on the spectral representation of ongoing EEG power, we observed the prototypical broad peak in the alpha frequency range (8 – 13 Hz; *Figure 2*). Moreover, alpha power decreased over the hemisphere contralateral to the attended stimulus position, indicating a functional disinhibition of cortical areas representing task-relevant regions of the visual field (Worden et al., 2000; Kelly et al., 2006; Thut et al., 2006). Concurrently, alpha power increased over the ipsilateral hemisphere, actively suppressing irrelevant and possibly distracting input (Rihs et al., 2007; Capilla et al., 2012).

A second approach focussed on the SSRs, i.e. strictly stimulus-locked rhythmic EEG components. As in classical frequency-tagging studies, we found spectrally distinct SSRs at the stimulation frequencies (here 10 and 12 Hz). These two concurrent rhythmic brain responses thus precisely reflected the temporal dynamics of the visual stimulation. Notably, SSR evoked power was between one to two orders of magnitude (10 – 100 times) lower than ongoing-alpha power. Smaller evoked power also explained why SSRs remained invisible in spectra of ongoing activity. They were likely masked by the broad alpha peak (Figure 2; Covic et al., 2017). Note that this is a result of the relatively low-intensity stimulation used here. Stimulation of higher intensity can evoke SSRs that are readily visible in power spectra of ongoing activity (Gulbinaite et al., 2019).

Crucially, we examined SSRs for effects of focused spatial attention. Visual cortical regions contralateral to the respective driving stimuli showed maximum SSR evoked power. We would expect to observe a decrease in SSR evoked power with attention (Kizuk and Mathewson, 2017; Gulbinaite et al., 2019) under the assumption that SSRs are frequency-specific neural signatures of a local entrainment of intrinsic alpha generators (Spaak et al., 2014; Notbohm et al., 2016) and exhibit similar functional characteristics. Instead, we found that SSR evoked power increased in line with earlier reports (Kim et al., 2007; Kashiwase et al., 2012; Keitel et al., 2013).

Note however that these attentional gain effects did not coincide topographically with scalp locations of maximum SSR evoked power (*Figure 4*). Instead, they were most pronounced over hemispheres ipsilateral to the position of the respective driving stimuli and thus co-localised with ipsilateral alpha power increases (*Figure 3*). Two control analyses showed that these effects were unlikely to be related (*Figure 5 & 6*). We have described the apparent counter-intuitive lateralisation of this effect before (Keitel et al., 2017a) when comparing scalp distributions by means of Attended-minus-Unattended contrasts (Keitel et al., 2017a). In that case, expecting attention effects to emerge at sites of maximum SSR power entails the implicit assumption that attention only acts as a local response gain mechanism. Alternatively, neural representations of attended stimuli could access higher order visual processing (Lithari et al., 2016) and a gain in spatial extent could then produce seemingly ipsilateral effects when evaluating topographical differences as observed here. However, previous cortical source reconstructions of SSRs in lateralised stimulus situations have unequivocally localised maximum effects of visuo-spatial attention to contralateral visual cortices (Müller et al., 1998b; Lauritzen et al., 2010; Keitel et al., 2013). Considering the limited spatial resolution of EEG, and that SSR inter-trial phase coherence showed yet another non-lateralised topographical distribution for gain effects (*Figure 4*), warrants a dedicated neuroimaging analysis of the underlying cortical sources that generate these attentional modulations.

### Opposite but co-occurring attention effects suggest interplay of distinct attention-related processes

Our analysis compared attention effects between “ongoing” spectral power within the alpha frequency band and a quantity termed SSR “evoked power” that is commonly used in frequency tagging research (Colon et al., 2012; Porcu et al., 2013; Stormer et al., 2014; Walter et al., 2016; Martinovic and Andersen, 2018). This term is somewhat misleading because it conflates a power estimate with the consistency of the phase of the SSR across trials of the experiment. Inter-trial phase consistency (ITC) is closely related to evoked power but involves an extra normalisation term that abolishes (or at least greatly attenuates) the power contribution^1^ (Cohen, 2014; Gross, 2014) and has been used to quantify SSRs before (Ruhnau et al., 2016).

The effects of attention on SSR evoked power and ITC are typically interchangeable (Covic et al., 2017; Keitel et al., 2017a). In fact, increased ITC, or phase synchronisation, has been considered the primary effect of attention on stimulus-driven periodic brain responses (Kim et al., 2007; Kranczioch, 2017). Looking at spectral power and ITC separately, as two distinct aspects of rhythmic brain activity, therefore resolves the attentional modulation conundrum: Seemingly opposing attention-related effects likely index different but parallel influences on cortical processing of rhythmic visual input. To avoid confusion, we therefore suggest opting for ITC (or related measures, e.g. the cosine similarity index (Chou and Hsu, 2018)) instead of “evoked power” to evaluate SSRs.

Incorporating our findings into an account that regards SSRs primarily as stimulus-driven entrainment of intrinsic alpha rhythms would require demonstrating how a decrease in alpha-band power (i.e. the contralateral alpha suppression) can co-occur with increased SSR phase synchronisation. Alternatively, stimulus-locked (“evoked”) and intrinsic alpha rhythms could be considered distinct processes (Freunberger et al., 2009; Sauseng, 2012). Consequentially, alpha range SSRs could predominantly reflect an early cortical mechanism for the tracking of fluctuations in stimulus-specific visual input per se (Keitel et al., 2017a) without the need to assume entrainment (Capilla et al., 2011; Keitel et al., 2014).

The underlying neural mechanism might similarly work for a range of rhythmic and quasi-rhythmic stimuli owing to the fact that visual cortex comprises a manifold of different feature detectors that closely mirror changes along the dimensions of colour, luminance, contrast, spatial frequency and more (Buracas et al., 1998; Blaser et al., 2000; Martinovic and Andersen, 2018). Most importantly, for (quasi-)rhythmic sensory input, attention to the driving stimulus may increase neural phase-locking to the stimulus to allow for enhanced tracking of its dynamics, i.e. increased fidelity. This effect has been observed for quasi-rhythmic low-frequency visual speech signals (Crosse et al., 2015; Park et al., 2016; Hauswald et al., 2018) and task-irrelevant visual stimuli at attended vs ignored spatial locations (Keitel et al., 2017a).

Concurrent retinotopic biasing of visual processing through alpha suppression and stronger neural phase-locking to attended stimuli could therefore be regarded as complimentary mechanisms. Both could act to facilitate the processing of behaviourally relevant visual input in parallel. In this context, SSRs would constitute a special case and easy-to-quantify periodic signature of early visual cortices tracking stimulus dynamics over time. Intrinsic alpha suppression instead may gate the access of sensory information to superordinate visual processing stages (Jensen and Mazaheri, 2010; Zumer et al., 2014) and enhanced ipsilateral alpha power may additionally attenuate irrelevant and possibly distracting stimuli at ignored locations (Capilla et al., 2012).

A neuronal implementation may work like this: During rest or inattention, occipital neuronal populations synchronise with a strong internal, thalamo-cortical pacemaker (alpha). During attentive processing of sensory input, retinotopic alpha suppression releases specific neuronal sub-populations from an internal reign and allows them to track the stimulus dynamics at attended locations. A related mechanism has been observed in the striatum, where local field potentials are dominated by synchronous oscillatory activity across large areas (Courtemanche et al., 2003). However, during task performance focal neuronal populations were found to disengage from this global synchronicity in a consistent and task-specific manner. At the level of EEG/MEG recordings, such a mechanism could lead to task-related decrease of oscillatory power but increase of coherence or ITC, as observed in the current study and previously in the sensorimotor system (Gross et al., 2005; Schoffelen et al., 2005; Schoffelen et al., 2011).

Whereas such an account challenges the occurrence of strictly stimulus-driven alpha entrainment, it may still allow alpha to exert temporally precise top-down influences during predictable and behaviourally relevant rhythmic stimulation – a process that itself could be subject to entrainment (Thut et al., 2011; Nobre et al., 2012; Haegens and Zion Golumbic, 2018; Zoefel et al., 2018).

### Conclusion

Our findings reconcile seemingly contradictory findings regarding spatial attention effects on alpha-rhythmic activity, assumed to be entrained by periodic visual stimulation, and SSRs. Focusing on spectral power or phase consistency of the EEG during visual stimulation yielded reversed attention effects in the same dataset. Our findings encourage a careful and consistent choice of measures of ongoing brain dynamics (here power) or measures of stimulus-related activity (here ITC), that should be critically informed by the experimental question, when studying the effects of visuo-spatial selective attention on the cortical processing of dynamic (quasi-) rhythmic visual stimulation. Again, we emphasise that both common data analysis approaches taken here can be equally valid and legitimate, yet they likely represent distinct neural phenomena. These can occur simultaneously, as in our case, and may index distinct cortical processes that work in concert to facilitate the processing of visual stimulation at attended locations.

## ACKNOWLEDGMENTS

Funded by a Wellcome Trust Joint Investigator Grant awarded to GT and JG (#098433/#098434). Lucy Dewhurst and Jennifer McAllister assisted in data collection. The experimental stimulation was realized using Cogent Graphics developed by John Romaya at the Laboratory of Neurobiology, Wellcome Department of Imaging Neuroscience, University College London (UCL).

## Notes

1 In a noisy, finite signal such as the typical second(s)-long EEG epoch, there will be a positive relationship between the power and inter-trial phase consistency at any frequency as is shown by the greater than zero noise floor in our ITC spectra (Figure 4). Also note that ITC only measures SSRs meaningfully if the neurophysiological signal contains a periodic component at the stimulation frequency.

## Author contributions

CK designed research, performed research, analysed data and wrote the article. JG designed research analysed data and wrote the article. AK, CSYB, CD and GT designed research and wrote the article.

## Data accessibility

EEG data, pre-processed in Fieldtrip format, that underlie all analyses reported here and a corresponding MATLAB analysis script are available on the Open Science Framework, osf.io/apsyf (Keitel et al., 2017b).

